# Direction-Averaged Diffusion-Weighted MRI Signal using different Axisymmetric B-tensor Encoding Schemes “Submitted to Magnetic Resonance in Medicine”

**DOI:** 10.1101/722421

**Authors:** Maryam Afzali, Santiago Aja-Fernández, Derek K Jones

## Abstract

**Purpose:** It has been shown previously that for the conventional Stejskal-Tanner pulsed gradient, or linear tensor encoding (LTE), as well as planar tensor encoding (PTE) and in tissue in which diffusion exhibits a ‘stick-like’ geometry, the diffusion-weighted MRI signal at extremely high b-values follows a power-law. Specifically, the signal decays as a 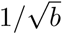 in LTE and 1/*b* in PTE. Here, the direction-averaged signal for arbitrary diffusion encoding waveforms is considered to establish whether power-law behaviors occur with other encoding wave-forms and for other (non-stick-like) diffusion geometries.

**Methods:** We consider the signal decay for high b-values for encoding geometries ranging from 2-dimensional planar tensor encoding (PTE), through isotropic or spherical tensor encoding (STE) to linear tensor encoding. When a power-law behavior was suggested, this was tested using in-silico simulations and in-vivo using an ultra-strong gradient (300 mT/m) Connectom scanner.

**Results:** The results show that using an axisymmetric b-tensor a power-law only exists for two scenarios: For stick-like geometries, (i) the already-discovered LTE case; and (ii) for pure planar encoding. In this latter case, to first order, the signal decays as 1/*b*. Our in-silico and in-vivo experiments confirm this 1/*b* relationship.

**Conclusion:** A complete analysis of the power-law dependencies of the diffusion-weighted signal at high b-values has been performed. Only two forms of encoding result in a power-law dependency, pure linear and pure planar tensor encoding and when the diffusion geometry is ‘stick-like’. The different exponents of these encodings could be used to provide independent validation of the presence of stick-like geometries in-vivo.

## 1 Introduction

Pathological disorders happen at the cellular level, therefore, it is important to obtain information on the micrometer scale. Diffusion MRI provides a tool to study brain microstructure based on the Brownian motion of water molecules [1] and it is therefore sensitive to the changes in the microstructure of the tissue [2, 3, 4]. Different mathematical representations are proposed to describe this relationship between the diffusion signal and the changes in the microstructure [5, 6, 7] the most prominent are the bi-exponential [8, 9, 10, 11, 12], the stretched exponential [13] and the power-law [14, 15, 16, 17]. The mathematical forms of these approaches are quite different. In the bi-exponential approach, the large b-value behavior is assumed to be dominated by the intracellular compartment. For stretched exponentials, the signal relationship with the b-value is exp [−(*kb*)^*a*^], where *k* is a constant and *a* < 1 is the stretching parameter. In the statistical model developed by Yablonskiy et al. [14], the signal decays as 1/*b* for large *b*, while the other studies [15, 16, 17], have reported that the signal at high b-value, decays as 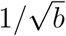.

The aforementioned studies all used single diffusion encoding. Since the development of the Pulsed Gradient Spin Echo sequence [18], there have been many works aimed at maximizing the information that can be obtained from a dMRI experiment by exploring different acquisition protocols [19, 20]. One such modification is the addition of multiple gradient pairs. We can use two pairs of pulsed-field gradients to obtain a Double Diffusion Encoding [21, 22]. It has been shown that double diffusion encoding, as well as other multiple encoding schemes such as triple diffusion encoding [23], provide information that is not accessible with single diffusion encoding [24].

This approach has been utilized by several groups for extracting microstructural information [25, 26, 27, 28, 29]. A framework was recently proposed [30] to probe tissue using different q-space trajectory encodings which can be described by a second-order b-tensor. Single, double and triple diffusion encoding can be characterized by b-tensors, with one, two and three non-zero eigen-values, respectively. In this framework, single diffusion encoding is also called Linear tensor encoding (LTE), double diffusion encoding with perpendicular directions is called planar tensor encoding (PTE) and triple diffusion encoding with equal eigenvalues is called spherical tensor encoding (STE). Herberthson et al. provided exact expressions for the direction-averaged signal obtained via general gradient waveforms [31]. They have shown that there is a power-law relationship for planar tensor encoding (*S* ∝ *b*^−1^).

In this study, we investigate the effect of different b-tensor encodings on the diffusion signal at high b-values. To remove the effect of fiber orientation distribution [32], the acquired signal is averaged over all diffusion directions for each shell. This so-called powder-averaged signal [33, 34] has less complexity than the direction-dependent signal. Powder averaging yields a signal whose orientation-invariant aspects of diffusion are preserved but with an orientation distribution that mimics complete dispersion of anisotropic structures.

We studied the power-law for different b-tensor shapes: i) to use it as cross-validation for the stick-like geometry. There is some combination of environments that give a power-law scaling using for example LTE but not necessarily using PTE. ii) to find the range of b-values that we observe this power-law scaling, iii) to investigate if the signal amplitude that we have in that range of b-value (7000 < *b* < 10000*s/mm*^2^) is considerable compared to the noise.

We also, studied the effect of the number of directions on this power-law scaling, for LTE and PTE schemes. One application of this study is to utilize the power-law representation of the LTE and PTE together to disentangle the intra-axonal signal fraction and the intra-axonal diffusivity from each other. Working in different range of b-values we can filter out some compartments based on the speed of the decay.

## 2 Theory

In multi-dimensional diffusion MRI, the b-matrix is defined as an axisymmetric second order tensor, *B* = *b/*3(1 − *b*_Δ_)*I*_3_ + *bb*_Δ_**gg**^*T*^, where **g** is the diffusion gradient direction and the b-value, *b*, is defined as the trace of the b-matrix. The eigenvalues of the b-matrix are 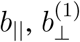 and 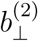 where 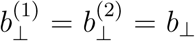 and *b*_‖_ is the largest. *b*_Δ_ is defined as *b*_Δ_ = (*b*_‖_ − *b*_⊥_)/(*b*_‖_ + 2*b*_⊥_). Changing *b*_Δ_, we can generate different types of b-tensor encoding. For LTE, PTE, and STE, *b*_Δ_ = 1, −1/2, and 0 respectively [23].

For the powder-averaged signal, the diffusion attenuation is a function of the orientation-invariant aspects of the diffusion and the encoding. The compartment diffusion attenuation is (Eq. (34) in [35]):

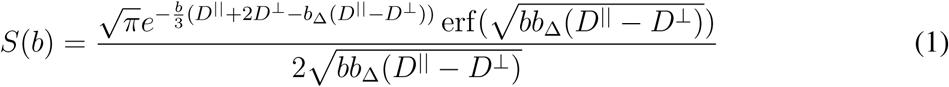

where *S* is the normalized diffusion signal and *D*^‖^ and *D*^⊥^ are the the parallel and perpendicular diffusivities respectively. We use the subscript “_*e*_” and “_*a*_” to denote the parameters of the extra- and intra-axonal compartments respectively.

Here, we study the effect of axisymmetric b-tensor shape on the diffusion-weighted signal at high b-values.

### 2.1 Linear, Planar and Spherical Tensor Encoding

In linear tensor encoding, *b*_Δ_ = 1 and assuming stick-like geometry, *D*^⊥^ = 0 in Eq. (1), therefore *S*_*ic*_ ∝ *b*^−1/2^. The sensitivity of MR to axon radius would alter the *b*^−1/2^ scaling [36] because there will be a perpendicular diffusivity and the exponential term in Eq. (1) will not be zero.

In planar tensor encoding, *b*_Δ_ = −1/2 and *S*_*ic*_ has the following form:

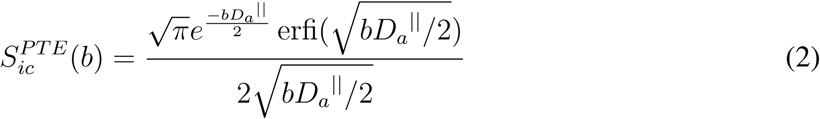

For large b-values, *bD*_*a*_^‖^ ≫ 1, therefore the diffusion signal can be approximated by the following equation (see Appendix A):

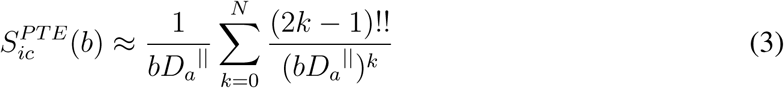

where !! denotes the double factorial and N depends on the *bD*_*a*_^‖^ value (Fig. 1 and Table 1.

**Table 1.** The minimum number of terms for reconstructing the PTE signal for different error threshold values.

| | | $bD_a^{\parallel}$ | | | | | | | | | | | | | | | | | |
| --- | --- | --- | --- | --- | --- | --- | --- | --- | --- | --- | --- | --- | --- | --- | --- | --- | --- | --- | --- |
|  |  | 3 | 4 | 5 | 6 | 7 | 8 | 9 | 10 | 11 | 12 | 13 | 14 | 15 | 16 | 17 | 18 | 19 | 20 |
| Error<br>threshold | 0.06 | 1 | 1 | 1 | 1 | 1 | 1 | 1 | 1 | 1 | 1 | 1 | 1 | 1 | 1 | 1 | 1 | 1 | 1 |
|  | 0.05 | - | 1 | 1 | 2 | 2 | 1 | 1 | 1 | 1 | 1 | 1 | 1 | 1 | 1 | 1 | 1 | 1 | 1 |
|  | 0.04 | - | 1 | 2 | 2 | 2 | 2 | 2 | 1 | 1 | 1 | 1 | 1 | 1 | 1 | 1 | 1 | 1 | 1 |
|  | 0.03 | - | 1 | 2 | 2 | 2 | 2 | 2 | 2 | 1 | 1 | 1 | 1 | 1 | 1 | 1 | 1 | 1 | 1 |
|  | 0.02 | - | 1 | - | 2 | 2 | 2 | 2 | 2 | 2 | 2 | 2 | 1 | 1 | 1 | 1 | 1 | 1 | 1 |
|  | 0.01 | - | - | - | 2 | 3 | 3 | 3 | 3 | 3 | 3 | 2 | 2 | 2 | 2 | 2 | 1 | 1 | 1 |

**Figure 1.**
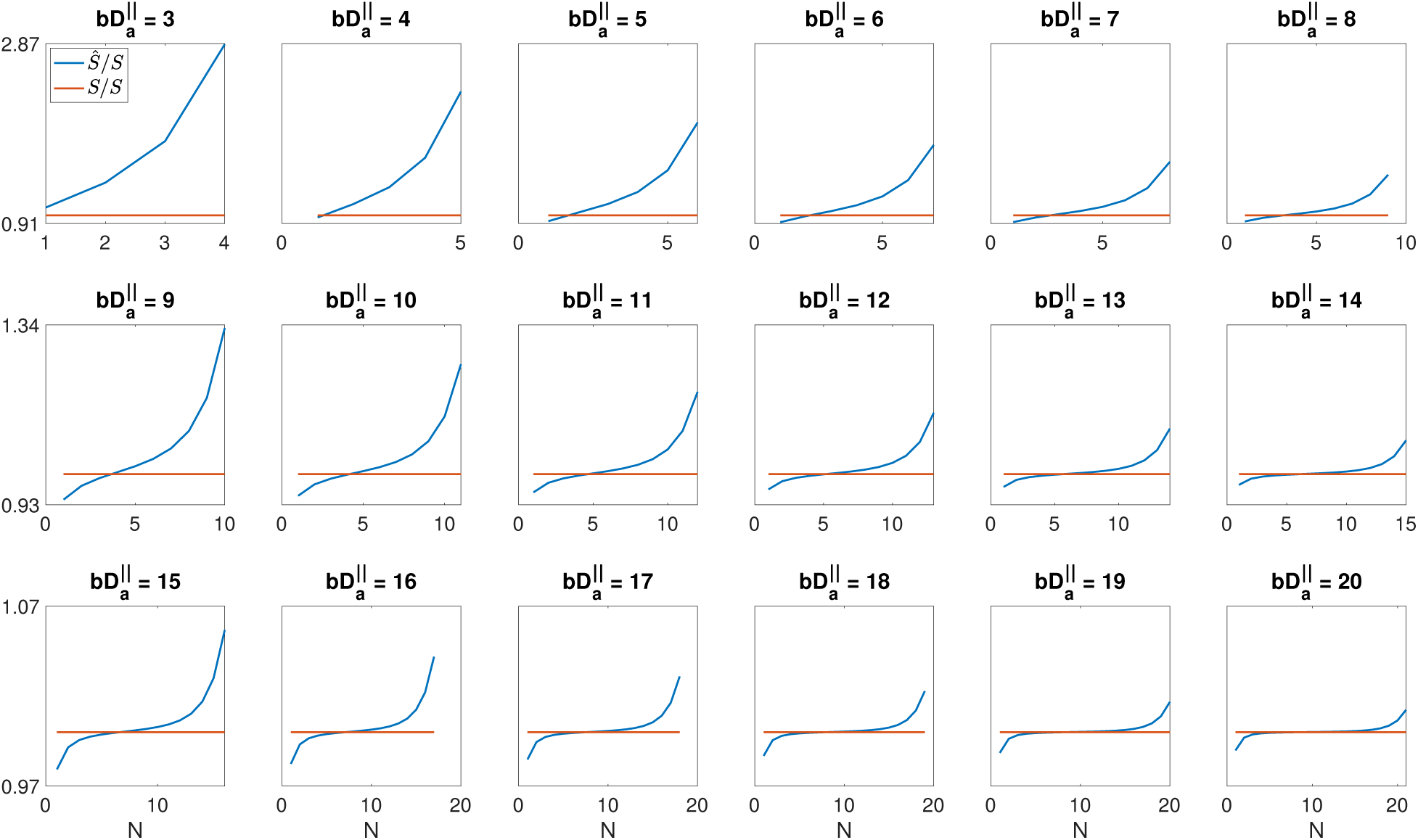
The approximated signal over the original PTE signal (*Ŝ/S*), for different *N* values.

For large b-values, the extra-axonal signal decays exponentially faster than the intra-axonal compartment, exp (−*bD*_*e*_^⊥^) ≪ 1, and can be neglected.

The asymptotic expansion of erfi(*x*) in Eq. 10 (see Appendix A) is valid when *x* → ∞, but large values of *bD*_*a*_^‖^ would suppress the signal to immeasurable levels, and therefore there are practical bounds on the value of *bD*_*a*_^‖^ that can be achieved. Therefore, we compared the original signal in Eq. 2 and the approximated signal using Eq. 3 for different values of *N* and *bD*_*a*_^‖^ (Fig. 1 and Table 1).We use a normalized error to compare the original (Eq. 2) and the approximated signal (Eq. 3):

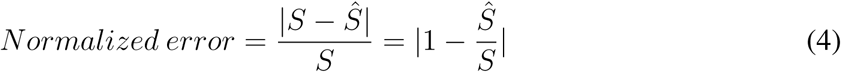

where *S* is the original signal obtained from Eq. 2 and *Ŝ* is the approximated signal from Eq. 3. In spherical tensor encoding, *b*_Δ_ = 0 and 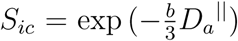. For large b-values, both intra- and extra-axonal signals decay exponentially fast, 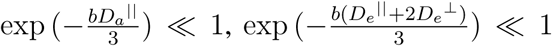 and both of them are negligible. Therefore, the spherical tensor encoding does not provide a considerable signal for large b-values in a two-compartment model.

### 2.2 General Case of Axisymmetric B-tensor

Here, we consider the general case of axisymmetric b-tensor *b*_Δ_ ≠ 0, to cover all b-tensor shapes between *b*_Δ_ = −0.5 (PTE) to *b*_Δ_ = 1 (LTE).

#### 2.2.1 0 < *b*_Δ_ ≤ 1

As noted above, in this range the error function in Eq. (1) goes to 1 for high b. To have a power-law relationship between the signal and the b-value, the exponential term exp[−*b*(*D*^‖^ + 2*D*^⊥^ − *b*_Δ_(*D*^‖^ − *D*^⊥^))/3] should go to one and therefore *D*^‖^ + 2*D*^⊥^ − *b*_Δ_(*D*^‖^ − *D*^⊥^) = 0. If *D*^‖^ = 0 then 2*D*^⊥^ + *b*_Δ_*D*^⊥^ = 0 and therefore *b*_Δ_ = −2 which is not in the range of plausible *b*_Δ_ values. For *D*^‖^ ≠ 0, *D*^⊥^/*D*^‖^ = (*b*_Δ_ − 1)/(*b*_Δ_ + 2) which is only physically plausible (i.e. the ratio of diffusion coefficients has to be ≥ 0) for *b*_Δ_ − 1 ≥ 0, but the maximum value that *b*_Δ_ can take is one, and therefore *D*^⊥^ has to be zero i.e. the geometry has to be that of a stick, and the b-tensor has to be a pure LTE to have a power-law relationship.

#### 2.2.2 −0.5 ≤ *b*_Δ_ < 0

Conversely, in the range −0.5 ≤ *b*_Δ_ < 0, as in Eq. (2), the error function becomes imaginary. Similar to the first scenario, to have a power-law relationship the exponential term has to be one. By replacing the first term of the approximation in Eq. (10) into Eq. (1), we have:

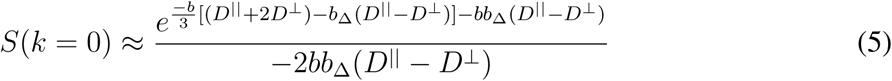

To have the exponential equal to one:

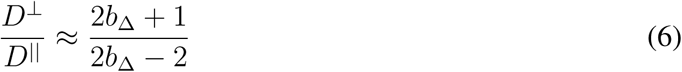

where the right side of the equation is negative for −0.5 < *b*_Δ_ < 0 which is not physically plausible for the left side of the equation (i.e. ratio of diffusivities). Therefore, the only possible case is to have *D*^⊥^ = 0 which again means stick-like geometry and *b*_Δ_ = −0.5 which is pure PTE. Clearly the exponential term will become zero if and only if *b*_Δ_ = −0.5, and thus the 1/*b* signal form will occur if and only if the b-tensor shape has just 2 non-zero eigenvalues, i.e. pure PTE. Thus, for stick-like geometries, there are only two b-tensor shapes for which a power-law exists: pure linear and pure planar. As the above equations show, we do not observe a power-law for non-stick-like geometries.

The *S* ∝ *b*^−1^ dependence is valid for an intermediate range of diffusion weightings and the asymptotic behaviour of the signal decay is determined by a steeper decay [37].

## 3 Method

### 3.1 Simulations

Synthetic data were generated with 60 diffusion encoding gradient orientations uniformly distributed on the unit sphere [38, 39] and 21 b-values spaced in the interval [0, 10000 *s/mm*^2^] with a step-size of 500 *s/mm*^2^. The noise is considered Rician with SNR = 150 for the b0 image, which is practically feasible using the Connectom scanner with an echo time of 88*ms* [40]. A three-compartment model with a Watson orientation distribution function is used:

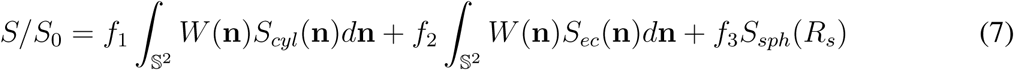

where *f*_1_, *f*_2_ and *f*_3_ are the intra-axonal, extra-axonal and the sphere signal fraction respectively, *W* (**n**) is the Watson ODF, *S*_*ec*_ is the extra-axonal signal, *S*_*cyl*_ is the signal attenuation of the impermeable cylinders [41] and *S*_*sph*_ is the restricted diffusion inside the spherical compartment in the presence of b-tensor encoding [42] (Appendix B). The ground truth parameter values defined by a set of parameters [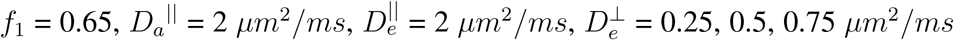, and *κ* = 11] and axon radius *r*_*i*_, come from the bins of the histograms in [43]. We average the signal over all *r*_*i*_s weighted by 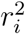. In histology, there is a possibility of tissue shrinkage. To account for this change, the axon radius values are multiplied with three shrinkage factors *η* = 0, 1, 1.5 [43, 44]. The *η* = 0 case simulates the effect of zero-radius axons.

The third compartment is simulated as a sphere with zero radius (dot) and a sphere with radius *R*_*s*_ = 8*µm* to consider the effect of combining the environments on the power-law scaling. The noisy diffusion signal is modeled according to the following:

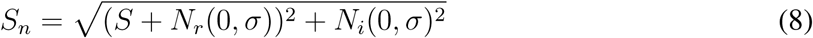

where *S*_*n*_ and *S* are the noisy and noise-free signal respectively and *N*_*r*_ and *N*_*i*_ are the normal distributed noise in the real and imaginary images respectively with a standard deviation of *σ* [45, 46]. The Matlab code for the simulation is available on GitHub (https://github.com/maryamafzali/PTE_Cylinder-)

### 3.2 In vivo Data

Two healthy participants who showed no evidence of a clinical neurologic condition were scanned in this study that was conducted with approval of the Cardiff University School of Medicine ethics committee. Diffusion-weighted images were acquired with 60 gradient directions for planar tensor encoding (PTE) on a 3T Connectom MR imaging system (Siemens Healthineers, Erlangen, Germany). Twenty axial slices with a voxel size of 4*mm* isotropic (given the strong signal attenuations investigated here, a low resolution of 4 mm isotropic was used) and a 64×64 matrix size, TE = 88 ms, TR = 3000 ms, were obtained for each individual.

Diffusion data were acquired for 10 b-value shells from 1000 to 10000 *s/mm*^2^ with a step size of 1000 and each shell had the same 60 diffusion encoding gradient orientations uniformly distributed on the unit sphere. One b0 image was acquired between each b-value shell as a reference.

The data were corrected for Gibbs ringing [47], eddy current distortions and subject motion [48]. To remove the Ricain noise ‘Non Local Spatial and Angular Matching’ method [49] was used. We normalized the direction-averaged signal based on the b0 signal in each voxel.

## 4 Results

Fig. 1 shows *Ŝ/S* for 3 < *bD*_*a*_^‖^ < 20 and 4 < *N* < 21. The selected range of *bD*_*a*_^‖^ is compatible with the range of b-values that we can obtain from the Connectom scanner and also the range of *D*_*a*_^‖^ that exist in the brain [50]. Based on Fig. 1 the number of terms in Eq. 7 should be smaller than or equal to the *bD*_*a*_^‖^ (*N* ≤ ⌊*bD*_*a*_^‖^ ⌋ where ⌊… ⌋ denotes the floor function) to have the minimum error (*Ŝ/S* is close to one). As the number of terms goes beyond the *bD*_*a*_^‖^, the error increases. Table 1 shows the minimum number of terms, *N*, for different error threshold values (0.01-0.06). When the error threshold is 0.02, we can approximate Eq. 2 with the first term in Eq. 3 if *bD*_*a*_^‖^ ≥ 14. For the error threshold of 0.06, the maximum *bD*_*a*_^‖^ to approximate the signal with the first term is 3.

Diffusion MRI is an inherently low SNR measurement technique, particularly when strong diffusion weightings are utilized. To reach the level that enables us to approximate the planar diffusion signal in Eq. 2 with the first term of Eq. 3, we need to use relatively high b-values (*bD*_*a*_^‖^ ≥ 14). One of the challenges with the high b-values is the noise, as the signal amplitude can be close to the noise floor. Therefore, here we find the maximum value of *bD*_*a*_^‖^ that we can use before hitting this rectified noise floor (see Appendix C).

The noise in complex MR data is normally distributed, whereas the noise in magnitude images is Rician distributed [45, 46]. Here, we select a minimum SNR value equal to 2 (see Appendix C). By setting the diffusion-weighted intensity to the mean background signal, we obtain the b-value that makes the signal equal to the noise floor.

Fig. 2 shows the maximum *bD*_*a*_^‖^ as a function of SNR for different encoding schemes and different noise floors. The maximum value of *bD*_*a*_^‖^ that can be used while staying above the noise floor increases when SNR increases, but the rate of this change is different for different encoding schemes. The maximum *bD*_*a*_^‖^ value 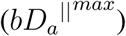 is proportional to the square of SNR, 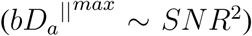 for LTE, where this relationship is linear for PTE (*bD*_*a*_^‖^ ∼ *SNR*) and it is logarithmic for STE (*bD*_*a*_^‖^ ∼ ln (*SNR*)). Based on this plot, if SNR = 50 the values of 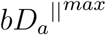 for linear, planar and spherical tensor encoding schemes are around 312, 21 and 9 respectively. The SNR in our data is around 250 [51] therefore the measured signal values in our experiment are higher than the noise level. For this SNR, the 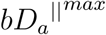 for linear, planar and spherical tensor encoding schemes are around 15625, 100 and 16 respectively.

**Figure 2.**
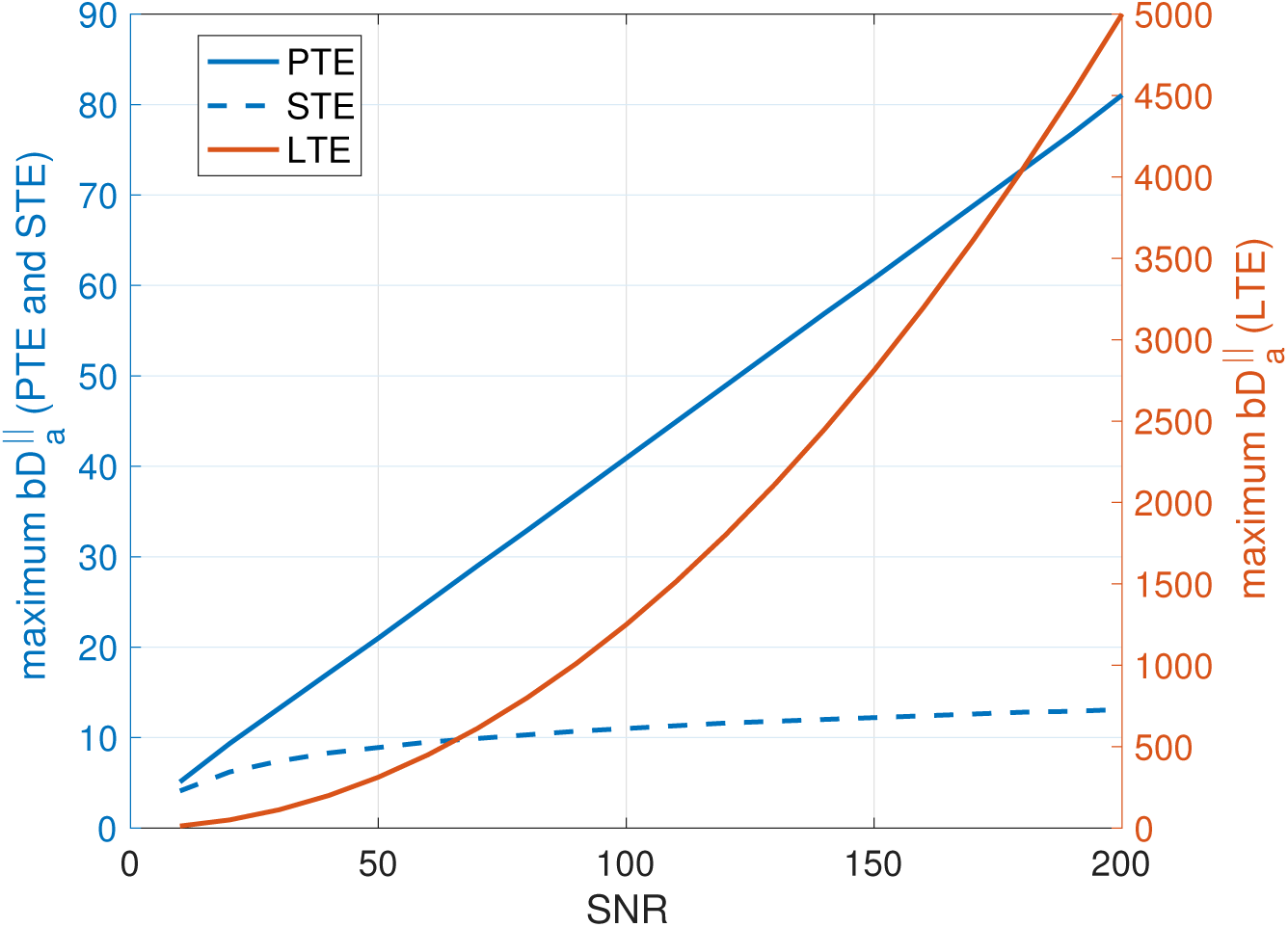
Maximum *bD*_*a*_^‖^ vs SNR. The Maximum *bD*_*a*_^‖^ value is proportional to the square of SNR, (*bD*_*a*_^‖^ ∼ *SNR*^2^) for LTE, where this relationship is linear for PTE (*bD*_*a*_^‖^ ∼ *SNR*) and it is logarithmic for STE (*bD*_*a*_^‖^ ∼ ln (*SNR*)).

Fig. 3 shows the simulated direction-averaged PTE signal (*f*_3_ = 0) as a function of 1/*b* for three different perpendicular diffusivities and three different shrinkage factors. The result of the power-law fit (*S* = *βb*^−*α*^) is represented by the red dashed line and the *α* and *β* values are reported in each plot. The trust-region-reflective algorithm is used for optimization with a fixed initial value (*α* = 1 and *β* = 0.2). The goodness of fit is evaluated using the Bayesian information criterion (BIC) [52], fully aware of how unreliable blind goodness-of-fit criteria is. In our simulation, *f* = 0.65, *D*_*a*_^‖^ = 2 *µm*^2^/*ms*, therefore if the approximation in Eq. 3 is valid, *β* ≈ 0.325 and *α* ≈ 1 indicate that the fit approximately matches the theory.

**Figure 3.**
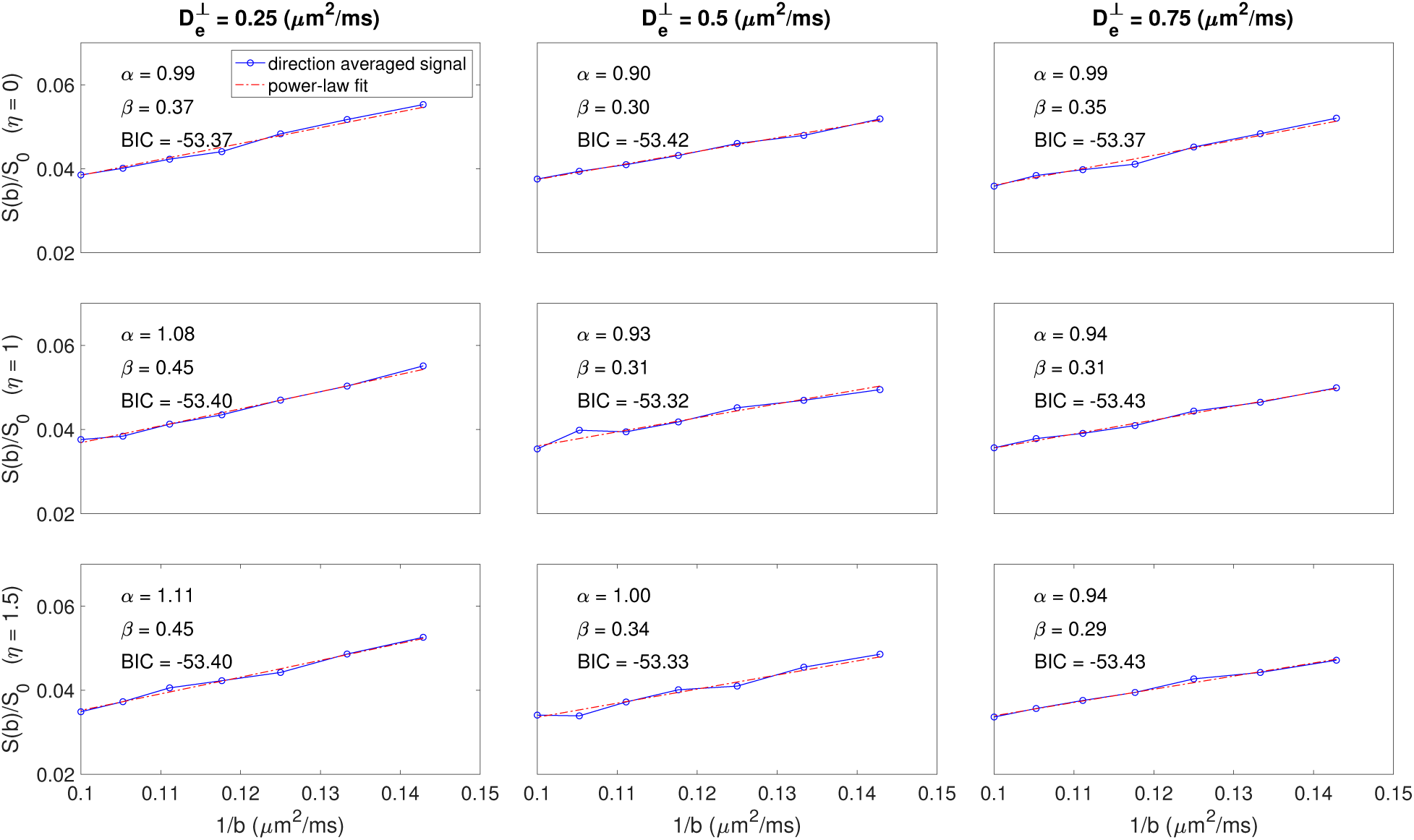
Simulated direction-averaged PTE signal for 7000 < *b* < 10000*s/mm*^2^ and the results of the power-law fit.

Szczepankiewicz et al. [53] showed that PTE needs less number of directions (15-20 directions for *b* <= 4000*s/mm*^2^) compared to LTE (20-32 directions *b* <= 4000*s/mm*^2^), to provide a rotationally invariant signal powder average, making it more efficient for powder averaging. Here we use a higher range of b-values 7000 < *b* < 10000*s/mm*^2^, therefore, the minimum number of directions for powder averaging is 45. Fig. 4 (a) shows the minimum number of directions for a rotationally invariant signal powder average for different b-values from 1000 to 10000 *s/mm*^2^ using LTE compared to PTE. Fig. 4 (b) illustrates the changes of exponent *α* using LTE compared to PTE. An insufficient number of diffusion directions in powder averaging may cause the break of power-law scaling. Therefore, we have to consider this when we use very high b-values.

**Figure 4.**
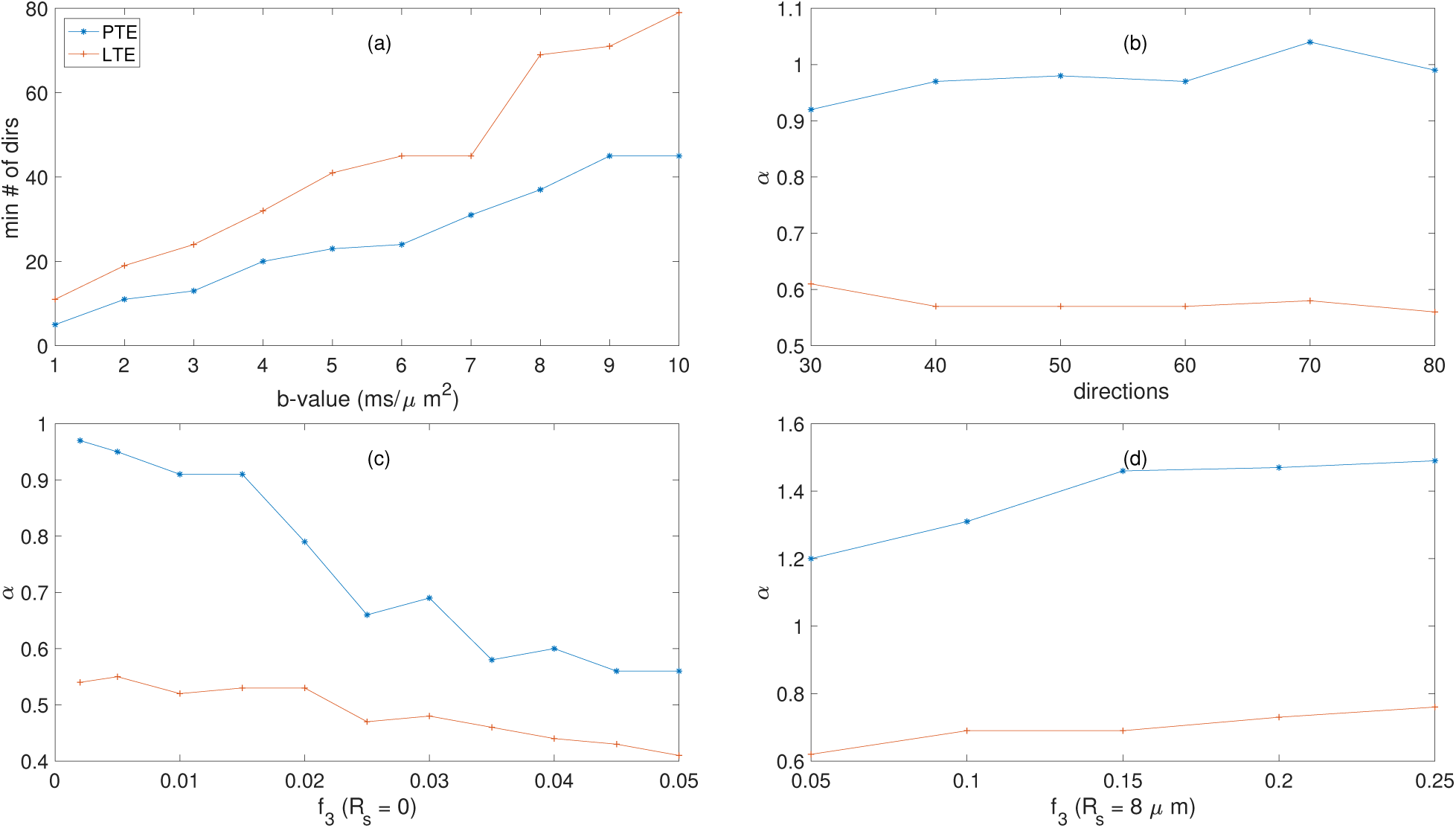
(a) The minimum number of directions for a rotationally invariant powder average signal in different b-values, (b) the changes of power-law scaling versus number of gradient directions, (c) the changes of the power-law scaling (*α*) versus ‘still water’ signal fraction, and (d) the changes of the power-law scaling (*α*) versus sphere signal fraction for PTE compared to LTE.

Still water compartment or ‘dot compartment’ has close to zero diffusivity. The presence of this compartment can affect power-law scaling. Fig. 4 (c) shows the changes of the power-law scaling (*α*) versus ‘still water’ signal fraction for PTE compared to LTE on the simulated data with *D*^⊥^ = 0.75(*µm*^2^/*ms*). Note that the *α* in PTE is more sensitive to the still water contribution than LTE.

Fig. 4 (d) shows the changes in the exponent *α* in the presence of spherical compartment with the radius of *R*_*s*_ = 8*µm*. The range of *α* values is close to the estimated values in gray matter (Fig. 5) which shows the combination of stick and sphere can represent the signal decay in gray matter [54]. Fig. 5 illustrates the normalized direction-averaged diffusion signal of the in vivo data for different b-values (7000 ≤ *b* ≤ 10000) in a few slices. For high b-values (7000 < *b* < 10000*s/mm*^2^) the amount of the signal is considerable compared to the noise and the white matter structure is completely clear in the images because of high SNR.

**Figure 5.**
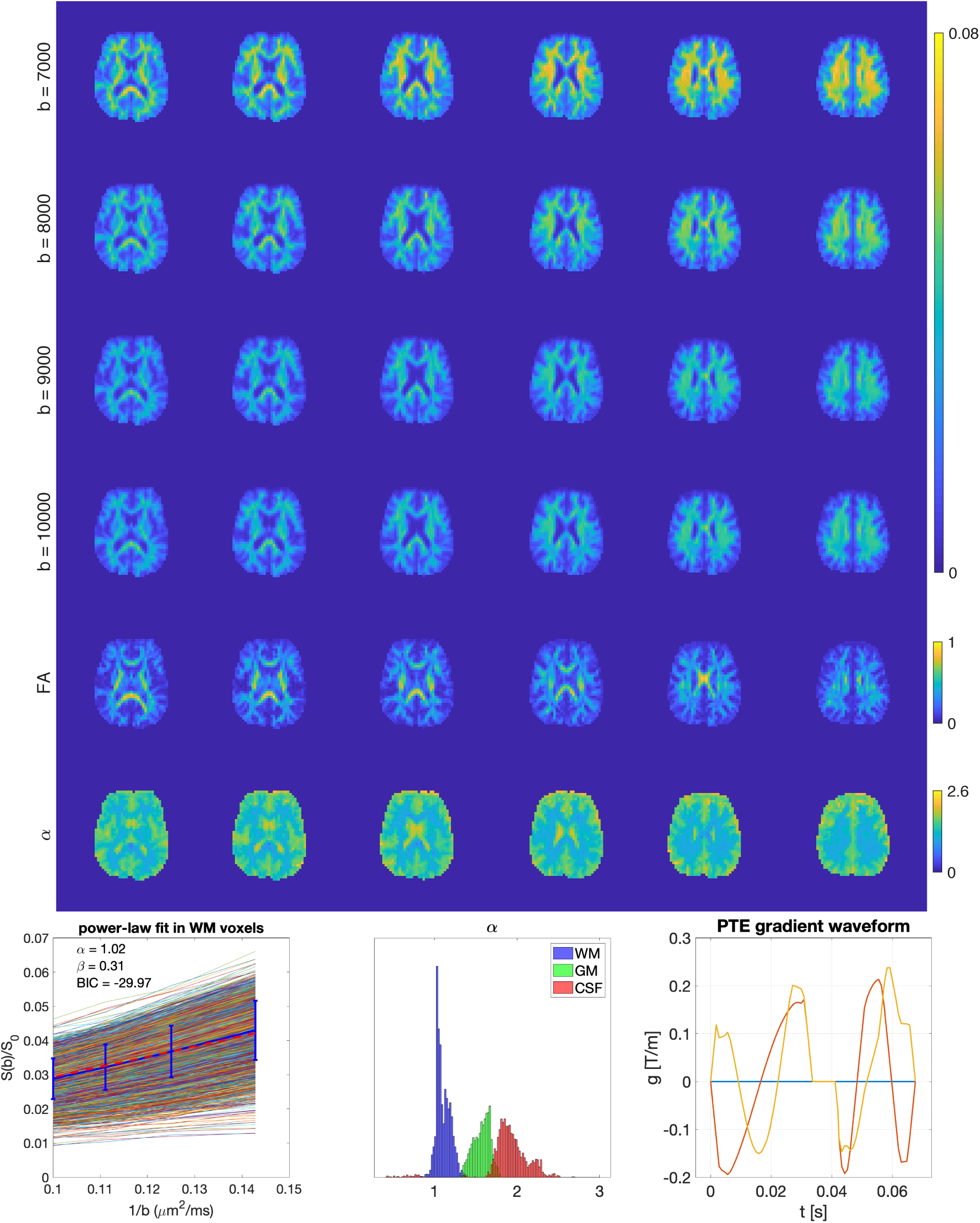
Direction-averaged diffusion signal for different b-values (b = 7000 to 10000 *s/mm*^2^) in PTE, FA, Parametric map of the exponent *α*. The plot of the diffusion signal vs 1/*b* for in vivo white matter voxels using planar tensor encoding. The blue curve with the error bar shows the mean and the std of the average signal and the red line shows the power-law fit. The parameters, *α* and *β* are reported in the figure. *α* = 1 shows the power-law relationship between the diffusion signal and the b-value. The histogram of *α* values and the PTE gradient waveform.

The last three rows of Fig. 5 show the FA and *α* map, the power-law fit over white matter voxels, the histogram of the *α* values in white matter, gray matter, CSF and the PTE gradient waveform. The PTE signal of the white matter voxels, the mean value, the standard deviation of the signal and the result of the power-law fit over the range of b-values investigated is shown in Fig. 5. The results show that the data are well described by power-law behavior, with *α* ≈ 1 which confirms the validity of the signal approximation using the first term in Eq. 3.

To segment the brain image into different tissues, we used FAST (FMRIB’s Automated Segmentation Tool) in FSL [55]. In the WM, the *α* value is close to one, supporting the theory. In grey matter and CSF, the exponent is larger (1.5 and 2, respectively). According to the theory outlined above, this would be consistent with a lack of pure ‘stick-like’ geometry in these tissue components. The spatial resolution of the data must be recognized, i.e., at 4 mm isotropic voxels, obtaining a ‘pure’ GM signal and ‘pure’ CSF signal is challenging. It is likely that the intermediate exponent in the GM between that of the WM and CSF is partly attributable to a partial volume effect, and partly attributable to the inadequacy of the model for grey matter architecture. The exponent in gray matter is similar to the one obtained using the combination of ‘stick + sphere’ [54]. Further investigation of this phenomenon in grey matter is beyond the scope of this work.

Table 2 shows the mean and the standard deviation of exponent *α* in white matter, gray matter and CSF for two different subjects. The mean value in WM is around one, in gray matter it is around 1.5 and for CSF around 2.

**Table 2.** The mean and the standard deviation of the exponent *α* in white matter, gray matter and CSF.

|  | WM | GM | CSF | BIC |
| --- | --- | --- | --- | --- |
| subject 1 | $1.1054 \pm 0.085$ | $1.5716 \pm 0.1009$ | $1.9004 \pm 0.2968$ | -29.9676 |
| subject 2 | $1.1100 \pm 0.1447$ | $1.6617 \pm 0.1836$ | $2.0469 \pm 0.2250$ | -30.293 |

## 5 Discussion

The main finding of this paper is a theoretical derivation, and confirmation in silico and in vivo of a power-law relationship between the direction-averaged DWI signal and the b-value using planar tensor encoding, as given by Eq. 3, for b-values ranging from 7000 to 10000 *s/mm*^2^. In white matter, the average value of the estimated exponent is around one.

For smaller b-values, this behavior must break down as the DWI signal of PTE cannot be approximated by Eq. 3 (Fig. 1) and also we cannot neglect the contribution of the extracellular compartment. It could also fail for very large b-values, if there were immobile protons that contributed a constant offset to the overall signal or if there is any sensitivity to the axon diameter [36]. Besides, if we do not have a sufficient number of diffusion directions for powder averaging, this power-law scaling can break.

The exponent of approximately one for white matter using PTE is consistent with the large b-value limit predicted for a model of water confined to sticks Eq. 3, which is used to describe the diffusion dynamics of intra-axonal water. Our results confirm this relationship between the diffusion signal and the b-value (Fig. 5 and table 2.

The *b*^−1/2^-scaling has previously been suggested by [17, 16] for linear tensor encoding. Two other proposed models predict power law signal decay, for large b-values using a linear tensor encoding. One of these is the statistical model [14], where the signal decays as 1/b for large b. Some other models [56, 57, 58], assume a gamma distribution for the diffusion coefficients and a family of Wishart distributions [59]. However, in this case, the exponent does not have a universal value, it depends on the distribution.

This work interprets the diffusion-weighted MRI signal decay at high b-values in the form of *S* ∼ *b*^−1^ for planar tensor encoding, this power-law relationship is also reported by Herberthson et al. [31]. An important application of this finding is using the combination of linear and planar tensor encodings to characterize the intra-axonal diffusivity and the signal fraction as it is proposed by [60] using triple diffusion encoding. PTE provides some information that is not available in STE. If we consider two different scenarios with the same mean diffusivity and different anisotropy where in the first one the perpendicular diffusivity is not zero while in the second one the perpendicular diffusivity is zero. Spherical tensor encoding gives the same signal for both scenarios while planar tensor encoding can distinguish these two from each other. Therefore, It is important to study the existence of power-law for planar tensor encoding which shows the presence of stick-like geometries. The amplitude of the signal that we get from STE for stick model in high b-values is less than the one that we get from PTE because the signal decay in STE is exponential while it is not exponential for PTE.

## 6 Conclusion

This work explores the diffusion-weighted MRI signal decay at high b-values for planar, and spherical tensor encoding complementing and extending previous works on linear tensor encoding. By exploring diffusion averaged signals, we conclude that the signal from STE decays exponentially for all the range of b-values. The intra-axonal signal does not decay exponentially as a function of b for linear and planar tensor encoding in high b-values. The direction-averaged DWI signal of PTE and LTE decreases with increasing b-values as a power law, for b-values ranging from 7000 to 10000 *s/mm*^2^. In white matter, the exponent characterizing this decrease is close to one-half, for LTE and one for PTE, which is consistent with the large b-value limit of a model in which intra-axonal water diffusion is confined to sticks. Obtaining an exponent of −1 for PTE and −1/2 for LTE could provide useful cross-validation of the presence of stick-like geometries in tissue. A complete analysis of the power-law dependencies of the diffusion-weighted signal at high b-values has been performed. Only two forms of encoding result in a power-law dependency, pure linear and pure planar tensor encoding. The different exponents of these encodings could be used to provide independent validation of the presence of stick-like geometries in vivo where, due to the slower decay, LTE is the most SNR efficient method. Any deviation from the power-law could indicate the deviation from stick-like geometry with both LTE and PTE encoding, as exploited by [36] for estimating the effective radius of axons. Again, for such applications of the power-law, the LTE approach is to be favored over the PTE approach, on account of the higher SNR per unit time.

## A Planar Tensor Encoding

In planar tensor encoding, *b*_Δ_ = −1/2 and *S*_*ic*_ has the following form:

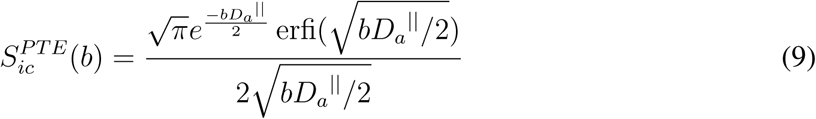

Asymptotic expansion of erfi(*x*) is as follows:

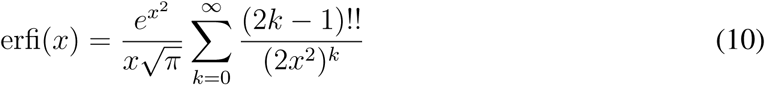

where *x* → ∞ and (−1)!! = 1.

*bD*_*a*_^‖^ ≫ 1 for large *b*, therefore we have:

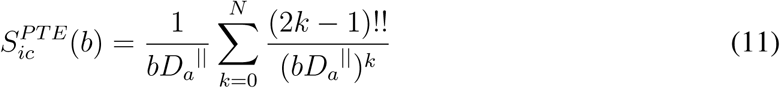

where N depends on the *bD*_*a*_^‖^ value (Fig. 1 and table 1.

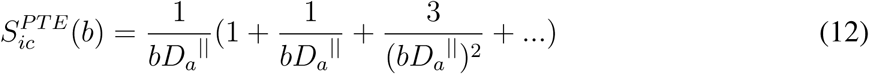

## B Signal Attenuation in a Cylindrical and Spherical Pore Using PTE

The signal attenuation of the impermeable cylinders [41] using PTE is generated using the following equation [42]:

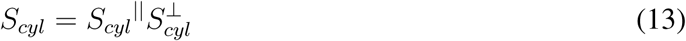

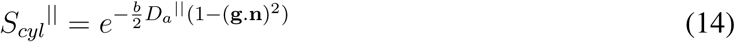

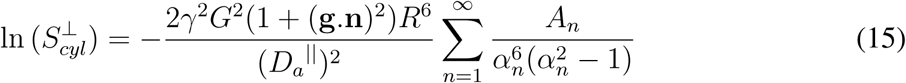

where *α*_*n*_ is the root of the derivatives of the first order Bessel function 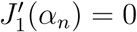 and

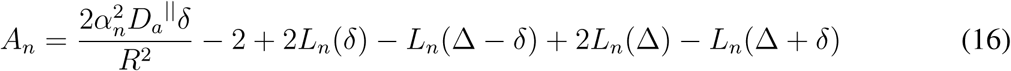

and

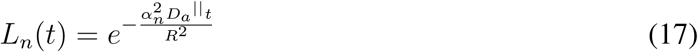

*S*_*Sph*_ is the restricted diffusion in spherical pore using planar tensor encoding [42]:

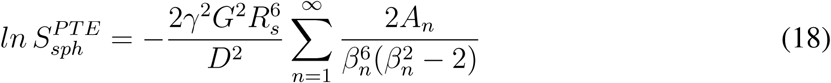

where *β*_*n*_ is the root of the derivatives of the first order spherical Bessel function 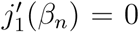. For the diffusivity of the water molecules inside the spherical pore (*D*), we use a constant value 1700*µm*^2^/*s*.

## C SNR and Error

Let us assume that a real signal *S* follows a Rician distribution with parameters *A* and *σ*

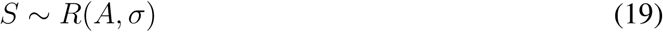

with PDF [45]

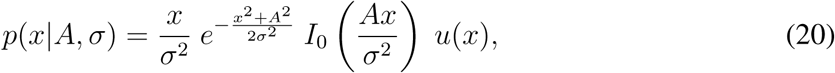

where *A* is the (absolute value) of the original signal (without noise) and *σ*^2^ is the variance of the complex Gaussian noise. It can be seen as

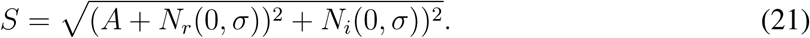

The question in MRI of how low can we go with the signal (i.e., when do we reach the noise floor) will always depend on the application and on the estimator we are using. However, we can always consider a lower bound to the SNR related to the error of the measured signal.

To calculate an SNR *threshold* independent of the particular application, we can use two different definitions of error: (1) the Mean Square Error (MSE) or (2) the mean error (ME). We define the MSE as

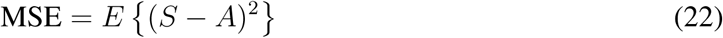

where *S* is the measured signal and *A* is the *original* signal. We use the mean value to assure that this error is a statistical property and not an isolated measure. The ME is alternatively define as:

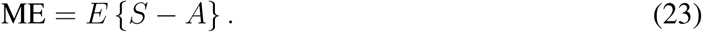

For the SNR calculation, we will consider that the error committed is a percentage of the original signal (to make it signal dependent), i.e.

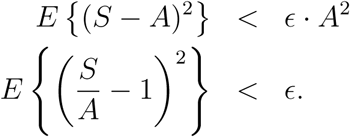

Alternatively, for the ME:

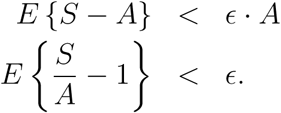

Assuming a Rician distribution of parameters *A* and *σ*, the errors become:

The MSE:

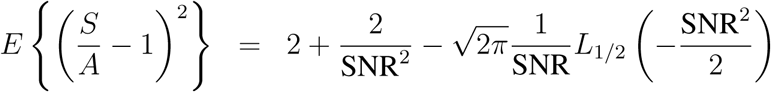

The ME:

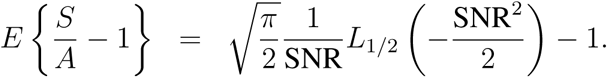

For the sake of simplicity, in this paper, we will consider ME as an error measure, since MSE is more restrictive. The relation between ME and SNR for different errors can be seen in Table 3.

**Table 3.** SNR values for different errors for the Rician model.

| Error | $< 0.005A$ | $< 0.05A$ | $0.1A$ | $0.2A$ | $< 0.5A$ |
| --- | --- | --- | --- | --- | --- |
| ME | SNR> 10 | SNR> 3.21 | SNR> 2.30 | SNR> 1.67 | SNR> 1.05 |

## Acknowledgements

The data were acquired at the UK National Facility for In Vivo MR Imaging of Human Tissue Microstructure funded by the EPSRC (grant EP/M029778/1), and The Wolfson Foundation. This work was supported by a Wellcome Trust Investigator Award (096646/Z/11/Z) and a Wellcome Trust Strategic Award (104943/Z/14/Z). S. Aja-Fernandez acknowledges the Ministerio de Ciencia e Innovación of Spain for research grants RTI2018-094569-B-I00 and PRX18/00253 (Estancias de profesores e investigadores senior en centros extranjeros). We are grateful to Emre Kopanoglu for feedback on the manuscript. We thank Zahra Moradi and Lars Mueller for help with the data acquisition. We thank Chantal Tax for help with the data acquisition and feedback on the manuscript. The authors would like to thank Filip Szczepankiewicz and Markus Nilsson for providing the pulse sequences for b-tensor encoding. We thank Marco Palombo for the fruitful discussion about the spherical compartment.

